# Partial decellularization of kidneys during prolonged acellular perfusion

**DOI:** 10.64898/2026.05.29.728736

**Authors:** Marlon J.A. de Haan, Annemarie M.A. de Graaf, Marten A. Engelse, Ton J. Rabelink

## Abstract

Long-term ex situ machine perfusion of donor organs is an emerging clinical strategy that creates a window for advanced therapeutic interventions. Replacing donor endothelial cells during machine perfusion with recipient endothelium could conceal the allogeneic epithelium from the recipient’s (humoral) immune response. We postulated that brief exposure to low concentrations of decellularizing agents could selectively remove vascular endothelium. Porcine kidneys were partially decellularized with five 2-minute infusions of either 0.01%, 0.1% or 1.0% SDS during acellular perfusion. As proof-of-concept, fluorescently-labelled porcine endothelial colony forming cells (ECFCs) were infused into the renal vein and artery of a partially decellularized kidney. Tissue analysis identified 0.1% SDS effectively removed endothelial cells from glomerular and peritubular capillaries. Infused ECFCs could be found back within the glomeruli. However, partial decellularization significantly impaired renal flow and increased vascular resistance. While partial decellularization successfully removed donor endothelium, the loss of vascular patency limits its clinical potential. Future research should prioritize modifying rather than removing donor endothelial cells.

## Introduction

Kidney transplantation is the only curative treatment for patients with end-stage renal disease (ESRD). While clinical outcomes have improved since the first transplants in 1954, recipients of allogeneic kidney grafts remain dependent on lifelong immunosuppression, which, despite increasing efficiency, imposes a cumulative lifelong burden. The ultimate goal, however, is to achieve graft tolerance and eliminate or reduce the need for these drugs.

Whole kidney engineering aims to achieve this by combining organ decellularization with subsequent repopulation using iPSC-derived nephron cell populations. Decellularization involves removing all cellular components by perfusing detergents, enzymes, or other cell-disrupting agents through the vascular network, while preserving the extracellular matrix (ECM), creating a decellularized kidney scaffold as a framework for reconstruction (1). Previously, we and others have demonstrated that the generation of such a scaffold is feasible, even for human kidneys (2, 3). However, generating and accurately delivering the necessary nephron cell populations to achieve functionality remains a significant challenge (4). For instance, while successful repopulation of vascular networks with (iPSC-derived) endothelial cells has been achieved in decellularized mouse, pig, and human kidney scaffolds – maintaining vascular patency in vivo up to seven days (3, 5) – a solution for adequate repopulation of the epithelial compartment has not yet been realized (4, 6).

Given these challenges, fully de- and recellularized kidneys are unlikely to address organ shortages in the foreseeable future. However, advances in decellularization and recellularization have paved the way for alternative approaches. One such strategy involves replacing a specific cellular compartment during ex vivo machine perfusion to improve donor-recipient compatibility. For highly sensitized patients, replacement of donor endothelial cells with recipient endothelium would conceal the underlying allogeneic epithelium from the recipient’s circulating immune system. Endothelial chimerism, combined with vascular sequestration of donor-specific antibodies, has been shown to prevent humoral rejection after islet transplantation (7).

However, such an approach would require a prolonged period of access to the organ. Over the past decade, machine perfusion has become a pivotal technology in organ preservation and transplantation (8, 9). Unlike static cold storage, which merely slows metabolic processes, dynamic machine perfusion supports organ metabolism, enables evaluation and monitoring of organ function, and provides a platform for advanced therapeutic interventions.

As the ability to support donor organs improves, machine perfusion is shifting from short-term use (hours) to extended durations (days). Recently, we established an acellular organ preservation platform that enables four-day ex vivo metabolic preservation of human kidneys (10). In this study, we assessed the potential of an engineering approach that involves partial decellularization of porcine kidneys during acellular perfusion and subsequent revascularization (**Figure 1**), with the aim of establishing endothelial chimerism in transplant organs. As a proof-of-concept, we infused porcine endothelial colony-forming cells into a partially decellularized porcine kidney to assess their ability to repopulate the vascular compartment.

**Figure 1.**
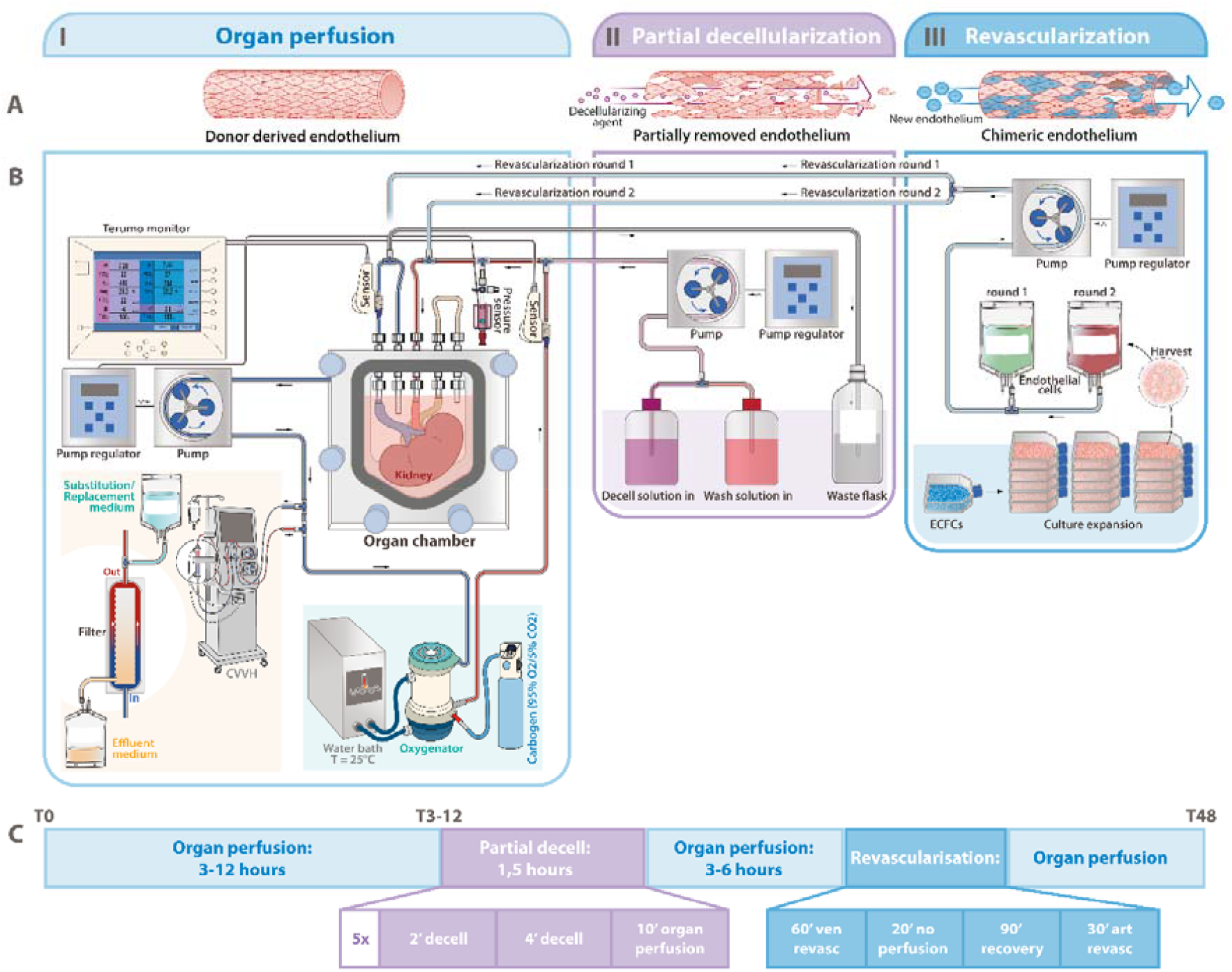
Conceptual and schematic overview for the partial decellularization and revascularization of kidneys during acellular perfusion. **A**, Conceptual overview of experimental approach, illustrating (I) the presence of donor-derived endothelium in the native kidney, (II) the partial decellularization of the donor organ vasculature achieved through brief periods of perfusion with decellularizing agents, and (III) subsequent revascularization using recipient derived endothelial cells. **B**, Organ perfusion platform and circuit components for partial decellularization and revascularization during multi-day organ perfusion. **C**, Experimental timeline demonstrating the sequence.

## Results

### Organ perfusion and partial decellularization platform

Abattoir-derived porcine kidneys were placed within a subnormothermic (25°C) acellular perfusion platform (**Figure 1A-B**). Renal flow was maintained by a pressure-controlled roller pump set at 75 mmHg, and kidneys were perfused for 3-12 hours before commencing partial decellularization. Partial decellularization was performed using five 2-minute infusions with a decellularizing agent, each followed by a 4-minute wash with DMEM F12 and a 10-minute perfusion with full perfusate between rounds (**Figure 1C**). We selected a range of decellularizing agents and concentrations – 0.01%, 0.1% and 1.0% sodium dodecyl sulfate (SDS), persufflation (air-perfusion) and trypsin (1x cell culture concentration) – to evaluate their effectiveness and minimize damage to non-target structures. For trypsin-based partial decellularization, the kidney underwent five cycles of 5-minute EDTA infusion, 5-minute trypsin infusion, and 5-minute PBS perfusion.

### Comparison of partial decellularization protocols

The efficacy of each protocol was assessed using periodic acid-schiff (PAS) staining, which highlights overall tissue and glomerular morphology, and Dolichos Biflorus Agglutinin (DBA) immunofluorescence, which stains endothelial cells in glomerular and peritubular capillaries (PTCs). Untreated and fully decellularized porcine kidneys served as positive and negative controls, respectively.

Histologic analysis demonstrated a dose-dependent effect of SDS on endothelial removal (**Figure 2**). Higher concentrations of SDS resulted in more extensive endothelial removal from both glomerular and peritubular capillaries, but also epithelial damage. While 0.01% and 0.1% SDS allowed degrees of endothelial cell removal without compromising epithelial structures, 1.0% SDS caused epithelial damage as reflected by gross morphology (**Figure 2**). Among the tested protocols, 0.1% SDS provided the best balance between endothelial removal and epithelial preservation (**Figure 2** and **Figure S1**).

**Figure 2.**
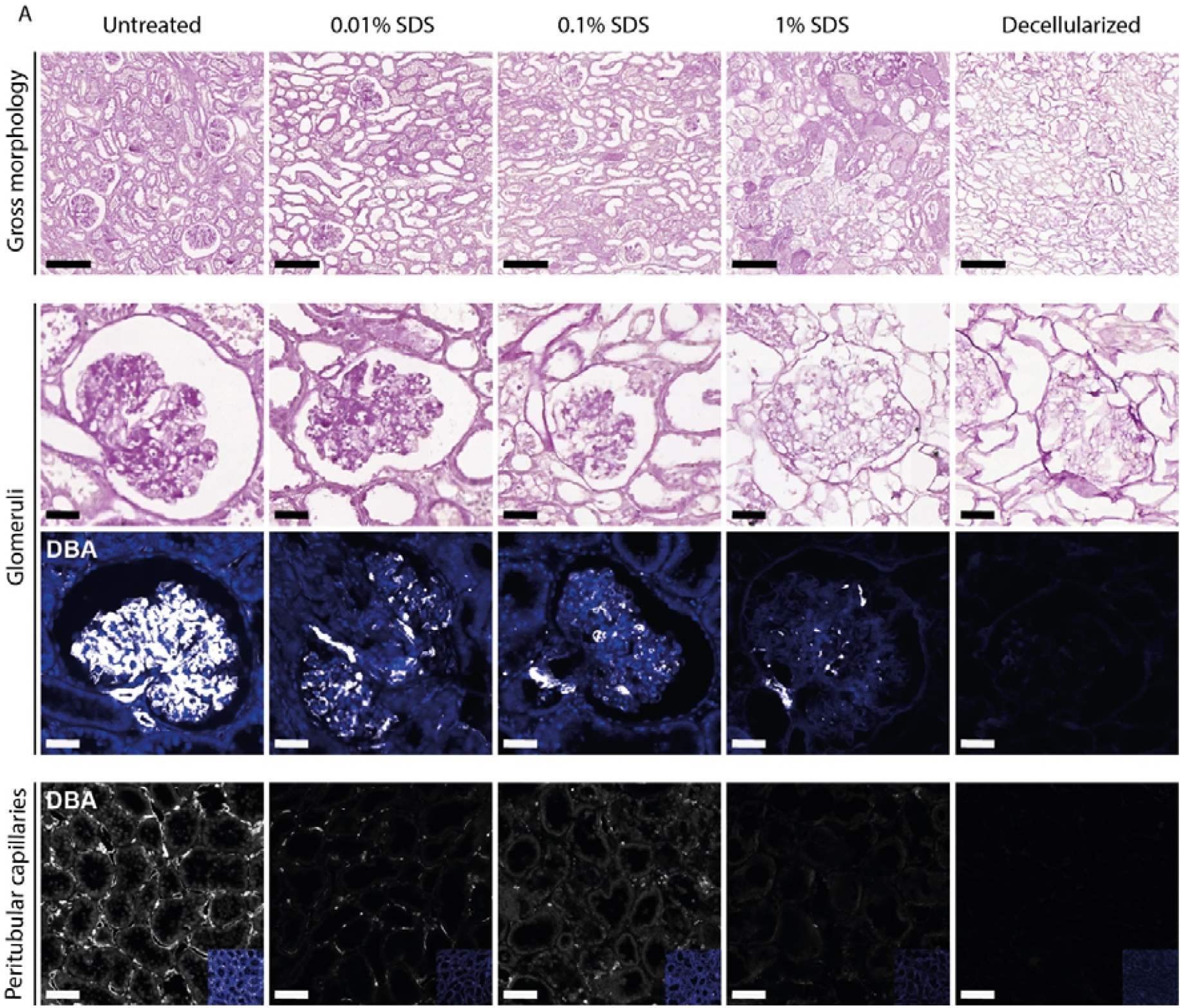
Optimizing partial decellularization protocols for donor endothelium removal. Histological assessment using Periodic acid Schiff (PAS) and Dolichos Biflorus Agglutinin (DBA) staining illustrates the effects on tissue structure (top row), glomeruli (middle rows) and peritubular capillaries (bottom row) across untreated, partially decellularized and decellularized porcine kidneys. Partial decellularization involved five 2-minute flushes with their respective SDS concentrations. Scale bars 100 μm for top and bottom rows, 40 μm for middle rows.

Persufflation and trypsin-based partial decellularization protocols were ineffective at removing endothelial cells from glomerular and peritubular capillaries (**Figure S2**). Additionally, trypsin-based protocols led to macroscopic structural disintegration in some of the perfused kidneys. Based on these findings, we selected five 2-minute infusions of 0.1% SDS as the most effective partial decellularization protocol.

### Proof-of-concept revascularization

Endothelial colony forming cells (ECFCs) from porcine peripheral blood were used for the revascularization of a partially decellularized kidney. Based on previous experiences (3), 200 million cells (100 million for each venous and arterial route) were labelled with a fluorescent label prior to infusion (venous infusion: green; arterial infusion: red).

Venous revascularization was initiated by switching from arterial to venous perfusion, and reducing perfusion pressure to 15 mmHg. Simultaneously, a -20 mmHg vacuum was applied within the organ chamber to enhance cell engraftment. Following venous cell infusion, renal flow was halted for 20 minutes before resuming arterial perfusion at 30 mmHg. After a brief stabilization period, arterial revascularization was performed. Thereafter, organ perfusion was continued for 18 hours for which perfusion pressure was increased to 70 mmHg.

Hemodynamic changes during partial decellularization and revascularization are visualized in **Figure 3A-C**. Of two paired and partially decellularized kidneys, the one with superior hemodynamic parameters was selected for revascularization, while the other was sampled for histologic assessment of partial decellularization (**Figure 3D**). In the revascularized kidney, arterially infused cells localized to the glomeruli (**Figure 3E**), whereas venously infused cells were not detected.

**Figure 3.**
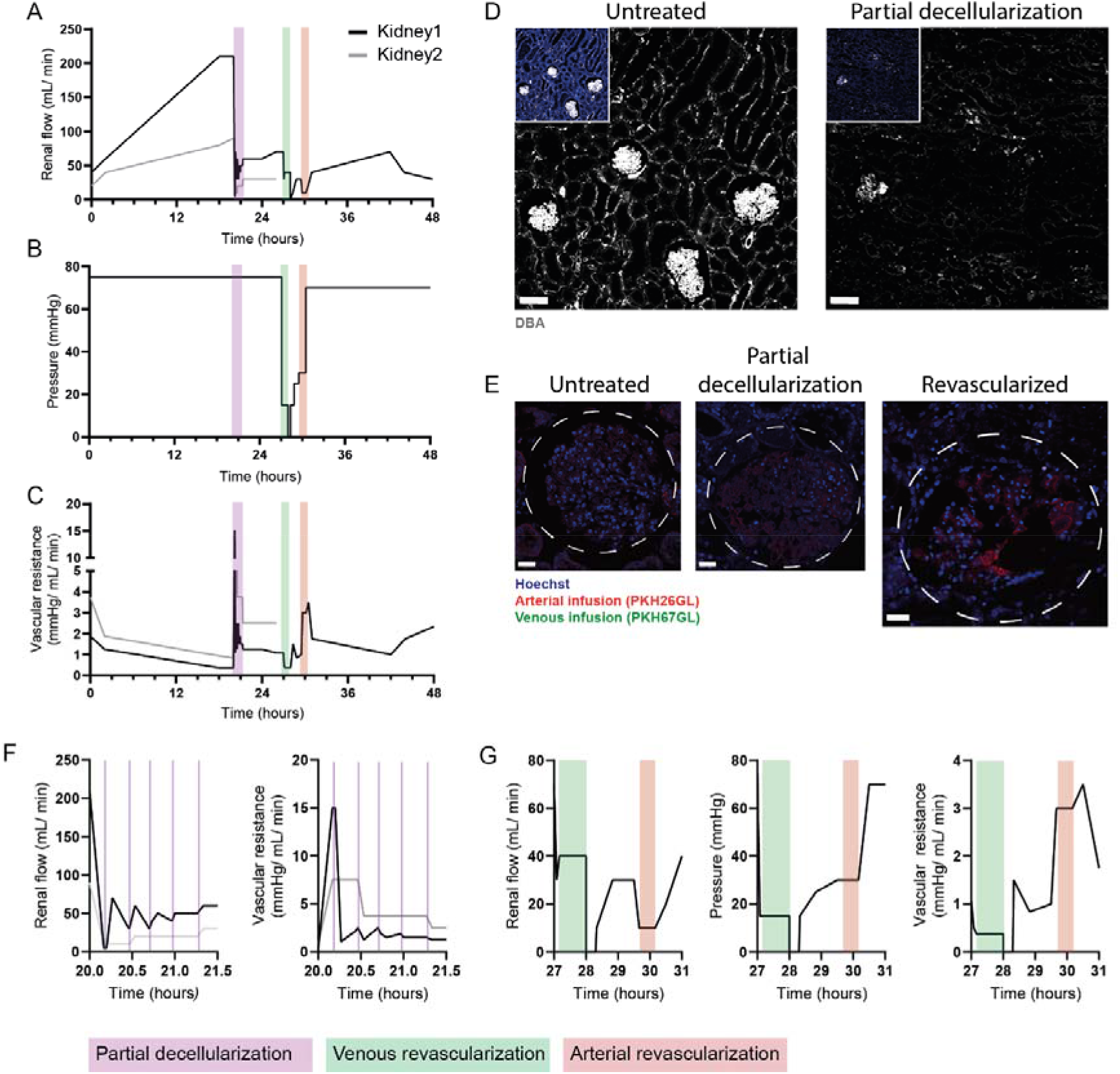
Proof-of-concept revascularization with endothelial colony-forming cells after partial decellularization. **A-C**, Perfusion hemodynamics during partial decellularization and subsequent revascularization. **D**, Comparison of Native and Devascularized porcine kidney. Scale bars 100 μm. **E**, Arterially infused ECFC were found back following revascularization within the glomeruli. Scale bars 40 μm. **F**, Detailed view of perfusion hemodynamics during partial decellularization. Vertical lines indicate the start of 0.1% SDS infusions. **G**, Detailed view of perfusion hemodynamics during venous and arterial revascularization.

### Bottlenecks

Exposure to SDS during partial decellularization led to an immediate increase in vascular resistance, as was reflected by a reduced renal flow (**Figure 4A**). Although flow partially recovered after decellularization, a significant reduction in renal flow and increase in vascular resistance persisted in partially decellularized kidneys(**Figure 4B-C**).

**Figure 4.**
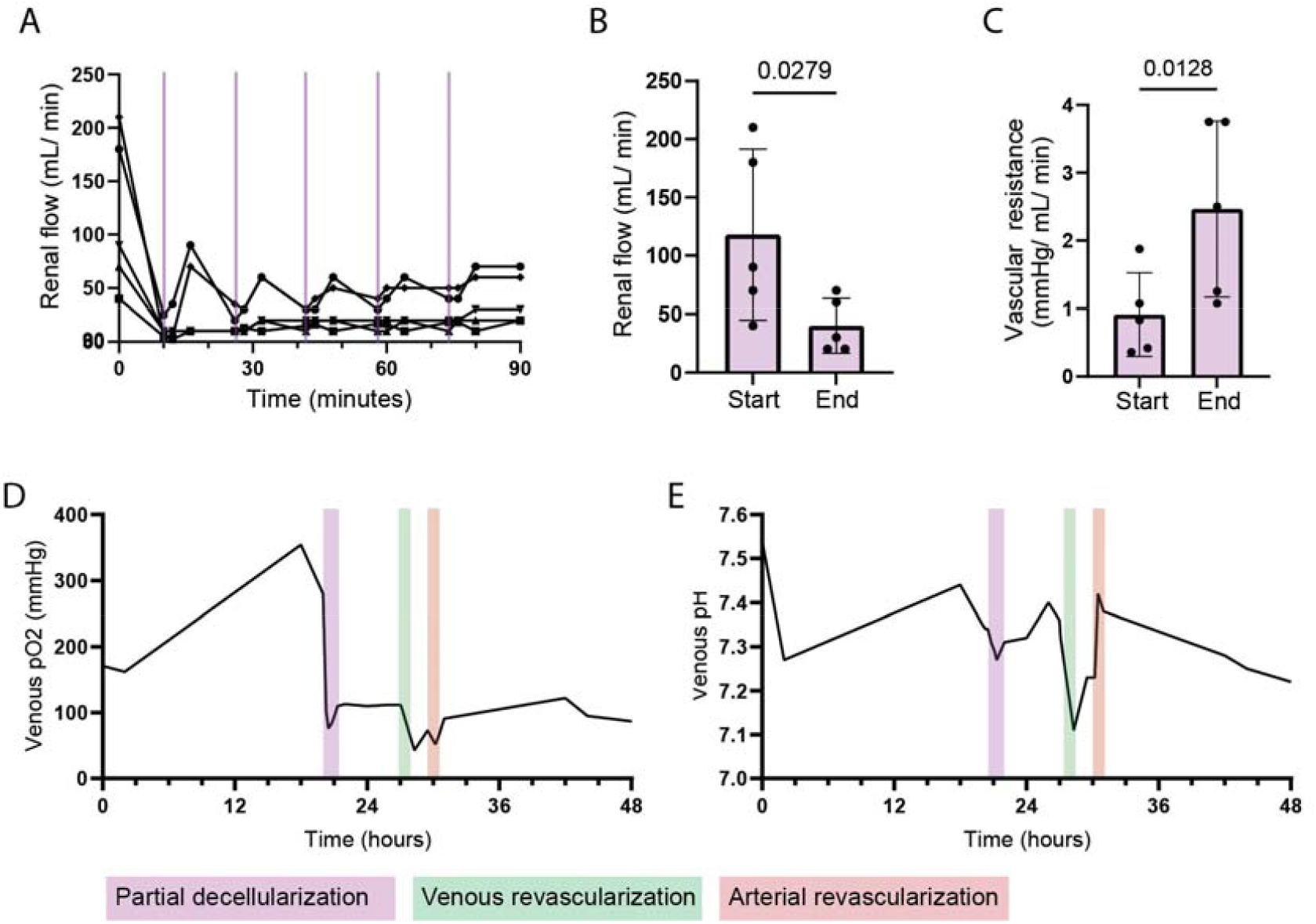
Key challenges during partial decellularization and revascularization. **A-C**, Hemodynamic changes during 0.1% SDS-based partial decellularization. Renal flow decreased significantly following partial decellularization (**B**), whereas vascular resistance increased (**C**). Paired t-test. **D-E**, Metabolic perfusate dynamics during 48-hour perfusion of a porcine kidney including partial decellularization and revascularization.

Additionally, venous pO_2_ and pH declined post-decellularization, indicating impaired metabolic function (**Figure 4D-E**). Metabolic deterioration continued to worsen after revascularization (**Figure 4D-E**).

## Discussion

In this study, we demonstrate that donor-derived endothelium can be selectively removed from kidneys during acellular perfusion, achieving an optimal balance between endothelial removal and epithelial preservation through five 2-minute infusions with 0.1% SDS. Theoretically, replacement of donor endothelium with recipient-derived or hypoimmunogenic endothelium could conceal the underlying allogeneic epithelium from the recipients circulating immune system, holding promise for highly sensitized patients.

Endothelial chimerism is a recognized phenomenon after organ transplantation (11), however, its biological significance remains debated. It is unclear whether chimerism reflects prior rejection episodes, is an adaptive mechanism by the recipient to align the graft with “self”, or a combination of both (12, 13). Unlike vascularized transplants such as the kidney, pancreatic islets are naturally revascularized by recipient-derived endothelial cells after transplantation (7). Notably, it has been demonstrated that endothelial chimerism, together with vascular sequestration of donor-specific antibodies, prevents humoral rejection. This points towards a potential immunological benefit of replacing donor endothelium within the transplant setting.

While partial decellularization removed the donor endothelium (**Figure 2**), it also led to increased vascular resistance (**Figure 4A-C**), which is concerning given the need to deliver oxygen and nutrients to sustain graft epithelium. Similar findings have been observed in rodent kidneys following partial decellularization (14). Towards the generation of endothelial chimerism, they infused placenta-derived endothelial cells into the renal artery during normothermic acellular perfusion, followed by 90 minutes of flow cessation to promote cell attachment, and subsequent perfusion for up to four hours.

To transfer revascularization methodology from rodent kidneys to human-scale organs we made several methodological adaptations. First, we infused ECFCs into both the renal vein and artery based on our experiences with whole kidney revascularization (3, 4). Second, perfusion was performed at subnormothermia (25°C), negating the need for oxygen carriers and enabling more extended perfusion durations (10). Finally, flow cessation was reduced from 90 to 20 minutes to balance epithelial preservation with endothelial attachment.

Despite these adaptations aimed to preserve the graft epithelium, our approach likely impaired ECFC engraftment, as indicated by the absence of detectable cells following venous infusion. For comparison, endothelial cells for whole kidney engineering are typically infused into fully decellularized scaffolds at 37°C, followed by up to three hours of perfusion cessation, also at 37°C, to maximize cell attachment and engraftment (3, 4). These conditions are simply incompatible with partially decellularized kidneys, where preservation of the epithelial compartment is essential. Our findings underscore the technical and logistical complexities of partial decellularization and revascularization of kidneys to achieve endothelial chimerism in human-scale transplant organs.

Cell replacement strategies during machine perfusion have been pioneered in other organs (15), such as the replacement of damaged airway epithelium in discarded human lungs (16, 17). In these studies, tracheal infusion of a decellularizing solution followed by epithelial cell replacement holds the advantage that the vasculature – and therewith supply of oxygen and nutrients – is not compromised. Similarly, cholangiocyte organoids have been demonstrated to repair bile ducts of the human liver during machine perfusion (18). These examples suggest that epithelial compartments may be more amenable to cell-based therapies during organ preservation.

In recent years, various innovative approaches have focused on (immuno)modulation of donor organs (19). These include nanoparticle-based targeting of the endothelium (20-22), enzymatic conversion of human blood group A organs to universal blood group O (23, 24), and cloaking of endothelial cells with immunosuppressive polymers (25). Notably, a recent breakthrough demonstrated the feasibility of silencing swine leukocyte antigen expression via lentiviral transduction of short hairpin RNAs, enabling porcine lung transplant survival without immunosuppression for up to 2 years (26). Of note, all these approaches rely on a “one-step” approach, in contrast to the “two-step” model of endothelial replacement explored in this study.

In conclusion, it seems improbable that replacing the endothelial lining within donor organs will have broad translational potential. Rather, future strategies should focus on modifying rather than replacing donor-derived cells. At this point, a future without further implementation and more widespread application of dynamic perfusion technologies for transplant organs seems unlikely. An ever increasing ability to support the needs of (injured) deceased donor organs allows a shift from short-term perfusion (hours) towards more extended durations (days). Therewith, a window of opportunity is created for the administration of more advanced therapies for graft (immuno)modulation, paving the way for a new era in organ transplantation.

## Methods

### Porcine kidney procurement

Kidneys were obtained from female landrace pigs (±6 months old, ±80 kg) from a local abattoir. Given the use of abattoir-derived porcine kidneys no animal ethical committee approval was required. Pigs were sedated with an electric shock followed by exsanguination and procurement, resulting in a warm ischemic time of 20-30 minutes. Kidneys were cannulated with Luer lock connectors (Cole Parmer) and flushed with 100-200 ml of cold Ringers acetate (B. Braun) supplemented with 12.500 IE L^-1^ heparin (LEO Pharma A/S), 10 mg L^-1^ butylscopolaminebromide (Boehringer Ingelheim) and 1 mg L^-1^ nitroglycerine (Hameln). Next, kidneys were flushed with 100 ml HTK (Histidine-Tryptophan-Ketoglutarate, Custodiol) preservation solution and transported on ice, resulting in a cold ischemic time of 3-6 hours.

### Acellular perfusion of porcine kidneys

Prior to perfusion, kidneys were flushed with 250 mL ice-cold DMEM F12 (Gibco). Acellular kidney perfusion was performed as previously described (10). Briefly, the platform was designed around an air tight organ chamber (Mascal Design). The renal artery, vein and ureter were connected to their designated outlets. A roller pump (Masterflex L/S Digital Drive 600 rpm) maintained pressure-controlled arterial perfusion at 75 mmHg. The acellular perfusate was oxygenated with a carbogen mixture of 95% O_2_ and 5% CO_2_ through a membrane oxygenator (Maquet Quadrox-I neonatal, Gettinge; or Sorin Lilliput 2, LivaNova) that was connected to a water bath. During the first hour of perfusion, temperature was increased from 10°C up to 25°C, and thereafter maintained at 25°C. Incorporation of a pediatric filter (Prismaflex HF20 set, Baxter) allowed the exchange of small molecular weight molecules. An in-line blood-gas sensor (CDI 500 system, Terumo Cardiovascular Systems) allowed monitoring of pO_2_, pCO_2_, pH and temperature in the venous outflow.

The acellular perfusate was composed of DMEM/F-12 (Gibco) was supplemented with Human Serum Albumin (HSA, Alburex 20, CSL Behring bv), Insulin-Transferrin-Sodium Selenite (ITS 100x, Sigma-Aldrich), sodium bicarbonate (Gibco), citric acid (Calbiochem, Merck), acetic acid (EMSURE, Merck), penicillin-streptomycin (Gibco), ciprofloxacin (Fresenius Kabi) and fungizone (Bristol-Myers Squibb). The total volume within the system was approximately 700 mL. Continuous hemofiltration was performed at 40 mL/h and filtration fraction of 5%. Substitution solution was added post-hemofilter by the Prismaflex system. The substitution solution contained the same components as the culture perfusate except for Human Serum Albumin

### Partial decellularization of porcine kidneys

After an initial perfusion phase (3-12 hours, depending on time of the day), various partial decellularization protocols were compared to assess the removal of donor endothelium within porcine kidneys. Partial decellularization was performed through five rounds of pressure-controlled (75 mmHg) 2-minute infusions with a decellularizing agent. The rounds were interspersed by a 4-minute wash with DMEM F12 and 10-minute perfusion with full perfusate. During the 2-minute infusion with a decellularizing agent and 4-minute wash all venous and urethral outflow was discarded.

We evaluated the effectiveness of different decellularizing agents, namely 0.01%, 0.1% and 1.0% sodium dodecyl sulfate (SDS), persufflation (air-perfusion) and trypsin (1x cell culture concentration). SDS was dissolved in DMEM F12 (Gibco). For trypsin-based partial decellularization, the kidney was treated with five rounds of 5-minute EDTA infusion, 5-minute trypsin infusion, and 5-minute PBS perfusion.

Untreated and fully decellularized porcine kidneys served as positive and negative control, respectively. Wedge biopsies were collected after partial decellularization for histological analysis. Whole porcine kidney decellularization was performed as previously described (3).

### Histological evaluation of partial decellularization

Wedge biopsies were fixed overnight in 4% paraformaldehyde, stored in 70% ethanol and embedded in paraffin for subsequent sectioning at 5 μm. For Periodic Acid-Schiff (PAS) staining, tissue sections were dewaxed, incubated for 30 min in 1% periodic water at 60°C and thereafter for 45 min in Schiff’s reagent at RT. Nuclear counterstaining was performed in hematoxylin solution. Sections were dehydrated and mounted with mounting medium (Entellan, VWR). For Dolichos Biflorus Agglutin (DBA) immunofluorescent staining, sections were dewaxed, antigens retrieved (Dako, s1699), incubated with a peroxidase blocking solution for 15 min (Abcam, ab64218), protein block solution for 60 min (Dako, X0909), incubated overnight at 4°C with 5 μg/ mL DBA in PBS (biotin-labelled; Sigma L6533), and incubated for 2 hours at RT with 4 μg/ mL Streptavidin A488 (ThermoFisher, S-11223). Nuclear counterstaining was performed with Hoechst. Sections were mounted with Prolong Gold (Invitrogen, P36930). All sections were digitized using a 3D Histech Pannoramic 250 Scanner (Sysmex) and viewed with CaseViewer software.

### Isolation of porcine endothelial colony forming cells

As cell source we opted for porcine peripheral blood-derived endothelial colony forming cells (ECFCs) given their accessibility, potential to be obtained from prospective transplant recipients and their scalability. ECFCs were isolated from heparinised porcine blood, as previously described (27). The peripheral blood mononuclear cell (PBMC) fraction was separated with Ficoll Paque (BD Biosciences), washed, and resuspended in EGM-10 medium (EGM-2 Basal Medium supplemented with growth factors (PromoCell), 10% heat-inactivated porcine serum, and 1% penicillin-streptomycin (Gibco)). Cells were seeded into rat tail collagen type I-coated (50 μg/mL; BD Biosciences) 48-well plates at a density of 250 000 to 400 000 cells per well. EGM-10 medium was refreshed every day for 7 days, and thereafter every other day. ECFC colonies started to appear from day7. Once colonies reached 80-90% confluence, they were collected and transferred to a 24 well. Colonies were pooled and culture expanded in T75 and thereafter T175 flasks.

### Revascularization of partially decellularized porcine kidneys with ECFCs

Prior to revascularization, 200 million ECFCs were labelled with PKH67GL or PKH26GL dyes for venous and arterial infusion, respectively. Labelled cells were resuspended in EGM-10 medium and transferred to infusion bags (Transfer pack, 600 mL, Fenwall Inc. [Baxter] Utrecht, the Netherlands) and placed on a shaker plate.

ECFCs were infused into both the renal vein and artery based on previous experiences with whole kidney revascularization (3). Venous revascularization was initiated by switching perfusion from the renal artery to the renal vein. Perfusion pressure was reduced to 15 mmHg, and a -20 mmHg negative pressure (vacuum) was created within the organ chamber by attaching a filter (Millex-FG, 0.20 μm, Merck) and LS-25 tubing (Masterflex) to the airtight organ chamber with a pressure probe (Edwards Lifesciences) and a custom-made pressure controller connected to a roller pump (Masterflex).

ECFCs were infused into the venous inflow at a rate of 4-8 mL/min over the course of 40 minutes. Following venous infusion, renal perfusion was stopped for 20 minutes before resuming pressure-controlled arterial perfusion at 30 mmHg. After a 60-minute stabilization period, arterial revascularization was performed by infusing cells at 4-8 mL/min into the arterial inflow. Perfusion was subsequently maintained for an additional 18 hours for which perfusion pressure was increased to 70 mmHg.

Wedge biopsies were taken at the end of perfusion and fixed in 4% paraformaldehyde or snap frozen in liquid nitrogen and stored at -80°C for subsequent analysis.

### Histological assessment of revascularization

Tissue biopsies were cryosectioned into 5-μm-thick sections using a Cryostar NX70 cryostat and mounted on glass cover slips. Nuclei were stained with Hoechst after which sections were imaged using SP8-WLL confocal microscopy and displayed using LAS-X software (Leica). Revascularization was assessed based on the presence or absence of PKH-dye labelled fluorescent cells.

### Statistical analysis

Data normality and equal variances were tested using the Shapiro–Wilk test. Statistical tests were performed using GraphPad Prism 9. P-values □≤□0.05 were considered statistically significant. Figures were created in Adobe Illustrator (Adobe Systems).

## Acknowledgments

This work is supported by the Dutch Kidney Foundation through the Participants of the Friends Lottery (20INI011). The Novo Nordisk Foundation Center for Stem Cell Medicine (reNEW) is supported by Novo Nordisk Foundation grants (NNF21CC0073729). Ton J. Rabelink is funded by the European Union through ERC grant (SPARK 101140863).

---

**Figure S1.**
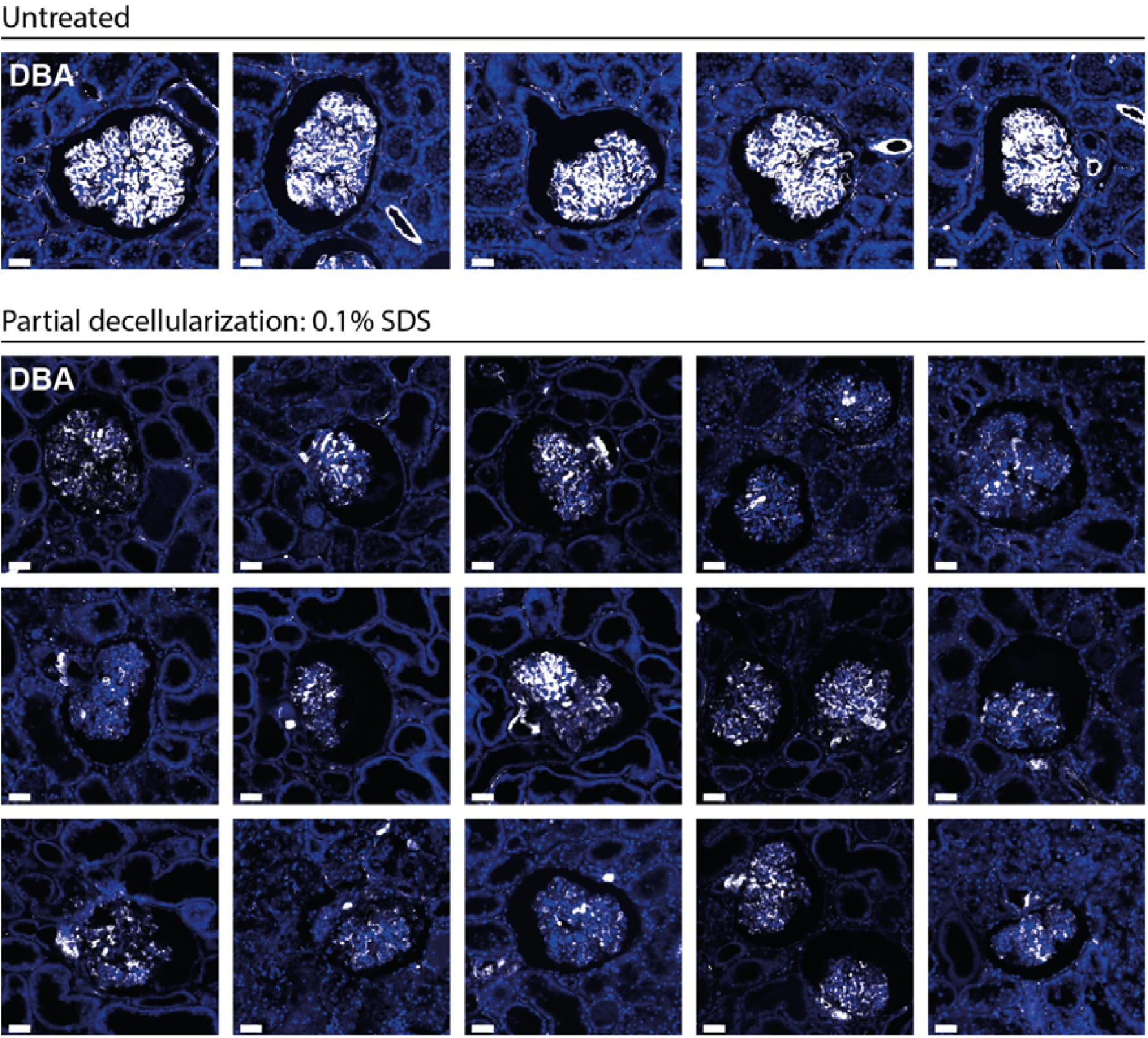
Glomerular vasculature after partial decellularization with 0.1% SDS. Histological assessment using Dolichos Biflorus Agglutinin (DBA) staining illustrates the partial decellularization of glomeruli following five 2-minute 0.1% SDS infusions. Scale bars represent 40 μm.

**Figure S2.**
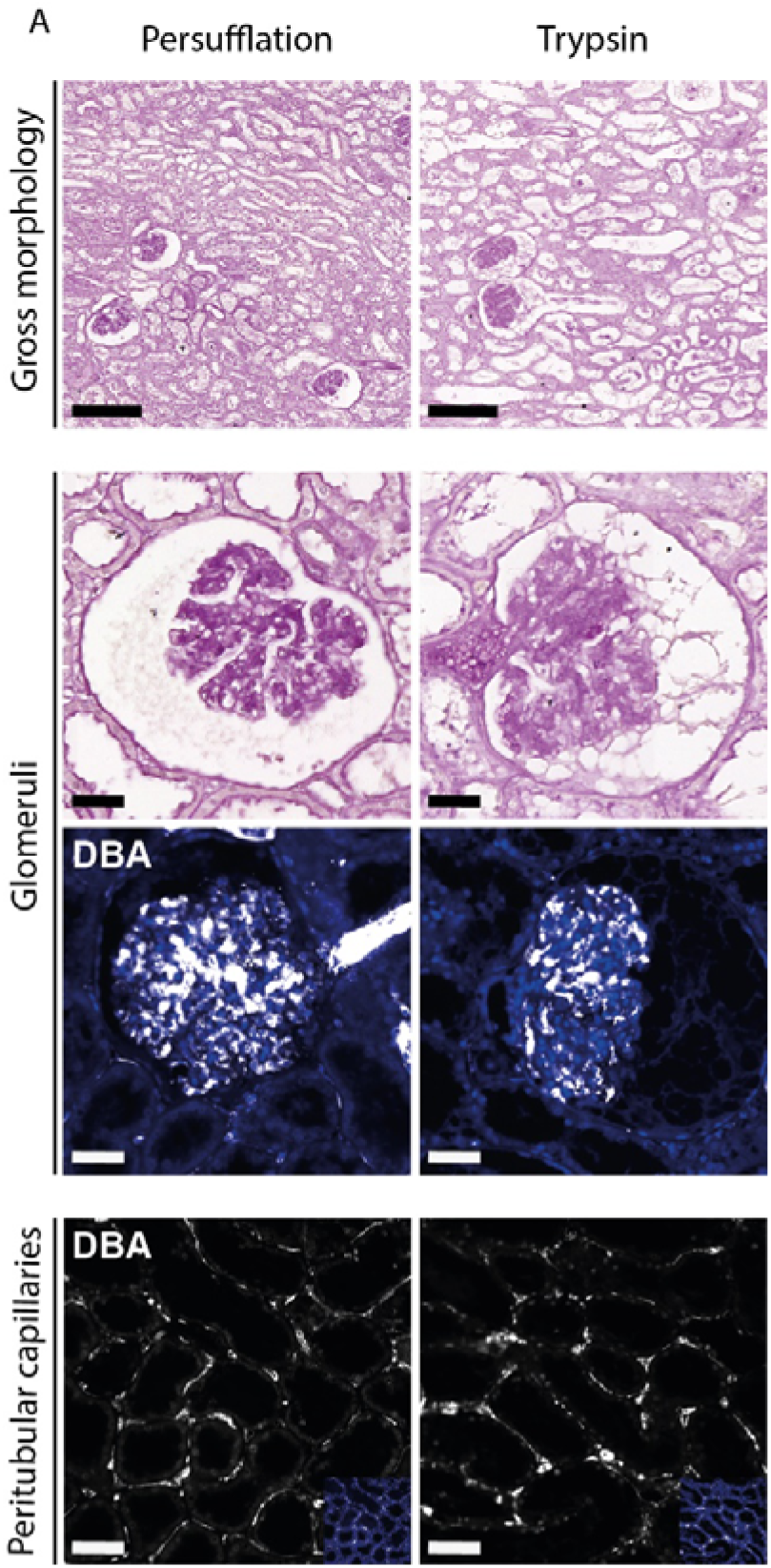
Partial decellularization using non-detergent-based protocols. Histological assessment using Periodic acid Schiff (PAS) and Dolichos Biflorus Agglutinin (DBA) staining illustrates the impact of two non-detergent-based partial decellularization protocols on tissue structure (top row), glomeruli (middle rows) and peritubular capillaries (bottom row). Persufflation-based partial decellularization entailed five cycles of 2-minute air perfusion, 4-minute DMEM F-12 perfusion, and 15-minute organ perfusion. Trypsin-based partial decellularization included five cycles of 5-minute EDTA perfusion, 5-minute trypsin perfusion, and 5-minute PBS perfusion. Scale bars 100 μm for top and bottom rows, 40 μm for middle rows.

## Notes

### Competing Interest Statement

The authors have declared no competing interest.

